# A 40 MHz femtosecond laser repetition frequency divider for enhanced two-photon imaging

**DOI:** 10.1101/2023.07.02.547378

**Authors:** Shiming Tang

**Affiliations:** Peking University School of Life Sciences, 100871 Beijing, China; Peking-Tsinghua Center for Life Sciences, 100871 Beijing, China; IDG/McGovern Institute for Brain Research at Peking University, 100871 Beijing, China; Key Laboratory of Machine Perception (Ministry of Education), Peking University, 100871 Beijing, China

## Abstract

Commercial Ti:Sapphire femtosecond lasers used for conventional two-photon microscopy typically operate at a ∼80 MHz repetition rate. However, this frequency is often suboptimal for cortical tissue imaging, where lower rates of 20–40 MHz are considered ideal. However, achieving these lower frequencies has remained a significant technical and financial challenge. Here, we present a compact, cost-effective resonant electro-optic modulator that halves the laser repetition rate to 40 MHz. When imaging neurons in the macaque visual cortex in vivo, this 40 MHz configuration yielded a >2-fold increase in fluorescence intensity and a ∼3 dB improvement in signal-to-noise ratio (SNR) for both green (GCaMP5G) and red (mScarlet) indicators. This enhancement proved particularly pronounced in deep-tissue imaging. Furthermore, the system demonstrated excellent long-term stability and induced no detectable phototoxicity. This simple and robust device represents a powerful upgrade for conventional two-photon microscopes, significantly enhancing imaging quality for the investigation of neural circuits within scattering brain tissue.

## Introduction

Two-photon laser scanning microscopy (2PLSM) has revolutionized biological imaging, enabling high-resolution, deep-tissue visualization of cellular and subcellular structures and dynamics within living organisms^1-3^. Its inherent optical sectioning capability and reduced scattering of near-infrared excitation light provide unparalleled access to the intricate workings of complex biological systems, particularly in the field of neuroscience for studying neural circuits in vivo^4-6^. The performance of 2PLSM, defined by imaging depth, signal-to-noise ratio (SNR), spatial resolution, and temporal resolution, is fundamentally constrained by the physics of two-photon excitation and the properties of the fluorescent probes and laser source employed^7,8^.

A critical parameter governing two-photon excitation efficiency is the laser pulse characteristic. The most widely used light source for 2PLSM is the mode-locked Ti:Sapphire laser, typically operating at a repetition rate of approximately 80 MHz dictated by their resonant cavity design. The quadratic dependence of two-photon absorption on instantaneous photon flux dictates that higher peak power per pulse yields exponentially greater fluorescence signal^9^. Consequently, for a fixed average laser power a crucial constraint to mitigate thermal damage and phototoxicity^10-12^ - reducing the laser repetition rate allows for an increase in the energy and peak power of individual pulses. This strategy has been shown to significantly enhance fluorescence excitation efficiency, thereby improving SNR and enabling greater imaging depth^13^. Previous studies have demonstrated that repetition rates in the range of 20-40 MHz can be optimal for deep-tissue imaging, balancing signal enhancement against increased risks of nonlinear photobleaching and photodamage associated with higher peak powers^14,15^. However, existing methods for reducing the repetition rate of standard 80 MHz lasers are often laden with compromises. While commercial pulse pickers and specialized low-repetition-rate laser systems are available, they can be prohibitively expensive^9^. Therefore, there remains a pressing need for a simple, cost-effective, and efficient method to halve the repetition rate of standard 80 MHz femtosecond lasers for widespread adoption in the neuroscience and biology communities.

Here, we present a compact and robust femtosecond laser repetition frequency divider that precisely converts an 80 MHz laser pulse train into a 40 MHz train. Our design utilizes a Pockels cell operating in a resonant model, driven by a custom-built, synchronized electronic module. This device can be seamlessly integrated into conventional 2PLSM setups. We demonstrate that, at the same average excitation power, our 40 MHz frequency divider achieves a two-fold increase in imaging intensity and a significant enhancement in SNR for both green and red fluorescent indicators in vivo, without introducing discernible phototoxicity in long-term imaging sessions. This simple yet powerful technical advance provides an accessible solution to enhance the performance of a vast number of existing two-photon microscopes.

## Results

### Design of a resonant 40 MHz frequency divider

To engineer the frequency divider, we first obtained a 40-MHz synchronizing signal by halving the 80 MHz clock from the femtosecond laser using an HMC432 frequency divider chip (**Fig. 1a, Supplementary Fig. 1**). For laser systems lacking a direct clock output, a photodiode-based circuit was used to extract the 80 MHz signal. The resulting low-amplitude (<1.0 Vpp) signal was then amplified to approximately 8 W using a 25 W RF power amplifier to drive a Pockels cell. A custom 1:10 impedance-matching transformer efficiently delivered this power to a resonant LC circuit built around the Pockels cell (**Supplementary Fig. 2**). The resonance was fine-tuned by adjusting the inductor coil; an 8-turn coil (8 mm diameter, 1 mm wire) was optimal for a Qioptiq LM 0202 P cell, whereas a 12-turn coil was required for a Conoptics 350-80-02 cell (**Supplementary Fig. 3a,b**). Precise phase alignment between the RF drive and the optical pulses, achieved by adjusting cable length, enabled the Pockels cell to function as a high-speed optical switch (Fig. 1b). This configuration deterministically routes alternating laser pulses into two distinct 40-MHz beam paths, effectively halving the repetition rate. For the Conoptics cell, active cooling was implemented to manage the high resonant current (**Supplementary Fig. 3b**).

**Fig. 1.**
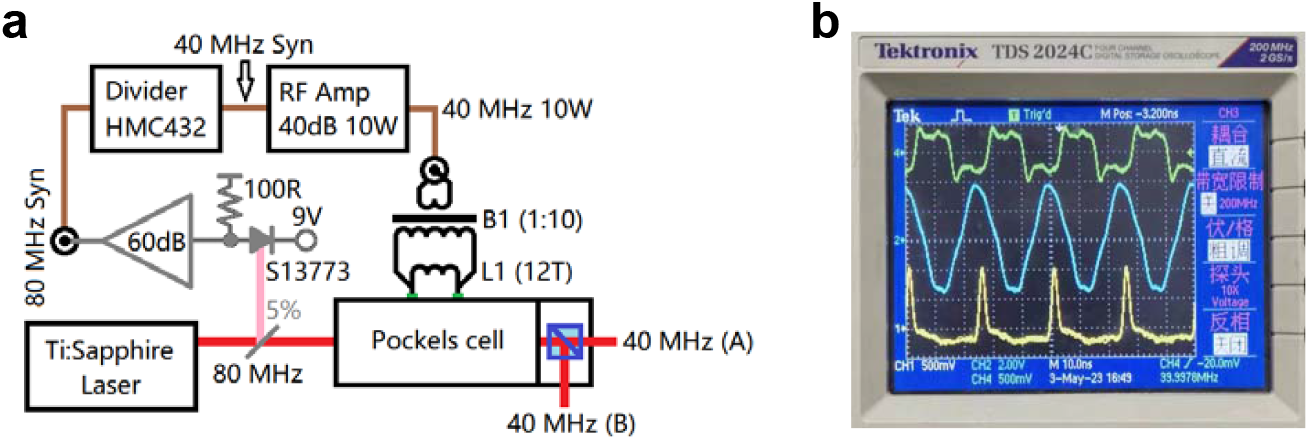
Design and signals of the 40 MHz femtosecond laser frequency divider. **a** Schematic of the resonant frequency divider. The device employs a Pockels cell operating in a 40 MHz resonant mode. The simplified driver circuitry consists of a frequency divider module, a 10 W commercial RF power amplifier, an impedance-matching transformer (B1), and an inductor (L1). **b** Oscilloscope traces of key signals. From top to bottom: the 40 MHz synchronization signal (green trace), the induced driving voltage from the resonant circuit (blue trace), and the output 40 MHz femtosecond pulse train as detected by an additional photodiode module.

### Reduced repetition rate enhances fluorescence signal and image quality

We first evaluated the performance of our system by conducting two-photon imaging of neurons expressing the calcium indicator GCaMP5G^6^ in the macaque visual cortex^16^. At a constant average power of 40 mW, switching from the native 80 MHz laser to the 40-MHz divided output produced a dramatic enhancement in image quality. We measured a greater than two-fold increase in mean fluorescence intensity and a corresponding signal-to-noise ratio (SNR) improvement of over 3 dB (**Fig. 2a, b**). Consequently, a 16-frame average acquired at 40 MHz surpassed the image quality of a 32-frame average at 80 MHz, effectively halving the required acquisition time for equivalent SNR (**Fig. 2c**). Notably, this enhancement did not compromise the temporal resolution required for resolving spontaneous calcium transients in neuronal somata (**Fig. 2d**).

**Fig. 2.**
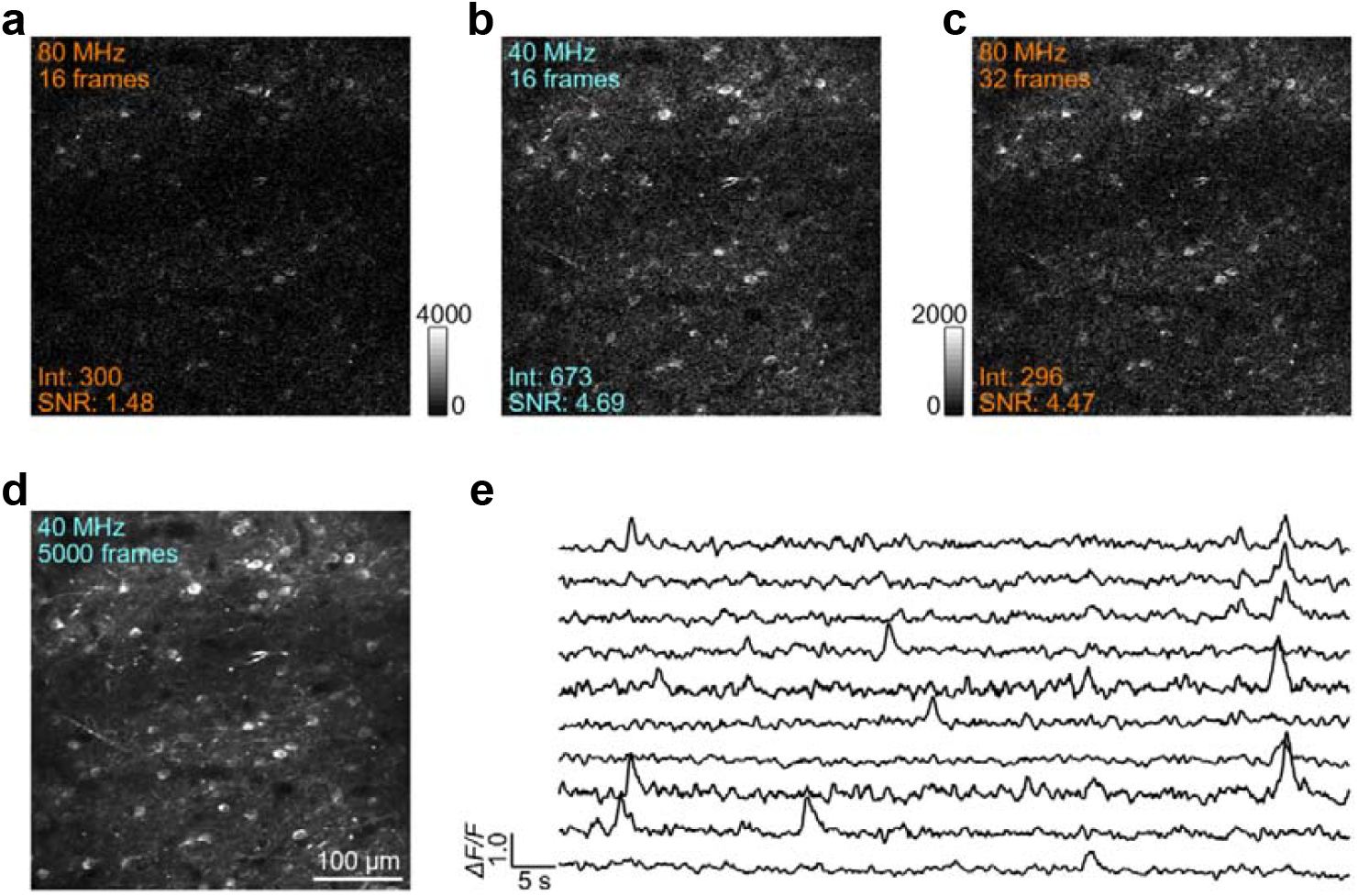
Comparison of two-photon imaging in macaque visual cortex using 80 MHz and 40 MHz laser excitation. **a** Two-photon image of cortical neurons acquired with a standard 80 MHz laser (16-frame average). **b** Image of the same field of view acquired with the 40 MHz frequency-divided laser at an identical average power of 40 mW (16-frame average). **c** Image from the 80 MHz acquisition, averaged over 32 frames, for quality comparison. **d** A long-exposure reference image from the 40 MHz recording, averaged over 5,000 frames. **e** Traces showing spontaneous calcium signals recorded from individual neurons within the field of view.

To confirm the broad applicability of our method, we repeated the experiment with neurons expressing the red fluorescent protein mScarlet3-H^17^. The 40 MHz excitation again conferred a substantial advantage, yielding a more than one-fold increase in fluorescence intensity and 2.4 dB SNR improvement at an average power of 48 mW (**Fig. 3**), demonstrating robust performance across different fluorophores.

**Fig. 3.**
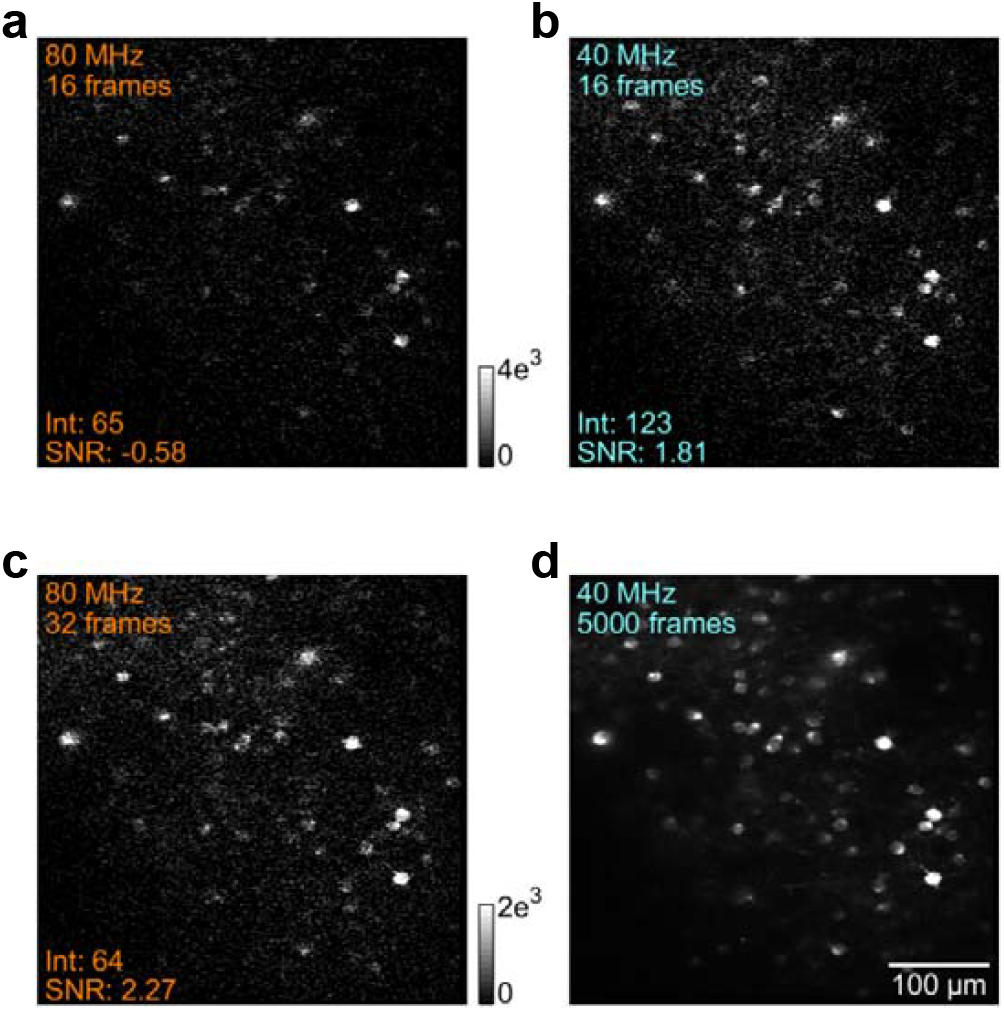
Enhanced two-photon imaging of macaque visual cortical neurons expressing the red fluorescent protein mScarlet3-H. All images were acquired at an average excitation power of 48 mW. **a** Image obtained with standard 80 MHz laser excitation (16-frame average). **b** Image obtained with the frequency-divided 40 MHz laser (16-frame average), showing significant improvements in signal intensity and structural clarity. **c** The 80 MHz acquisition averaged over 32 frames for image quality comparison. **d** A reference image generated by averaging 5,000 frames from the 40 MHz dataset.

### The 40 MHz mode enables superior deep-tissue imaging

The increased pulse energy of the 40 MHz laser train is particularly advantageous for imaging deep within scattering tissue. We tested this hypothesis by imaging neurons at a depth exceeding 600 μm in the macaque cortex. At this depth, the performance gain was even more pronounced than in superficial layers, with the 40 MHz mode yielding a 146% increase in image intensity and a 3.85 dB improvement in SNR (**Fig. 4a, d**). Under these challenging conditions, fine neuronal processes, such as a thin dendrite at 609 μm depth, were clearly resolved using the 40 MHz laser, whereas they were largely obscured by noise with the 80 MHz laser at the same average power (**Fig. 4c** and **f**).

**Fig. 4.**
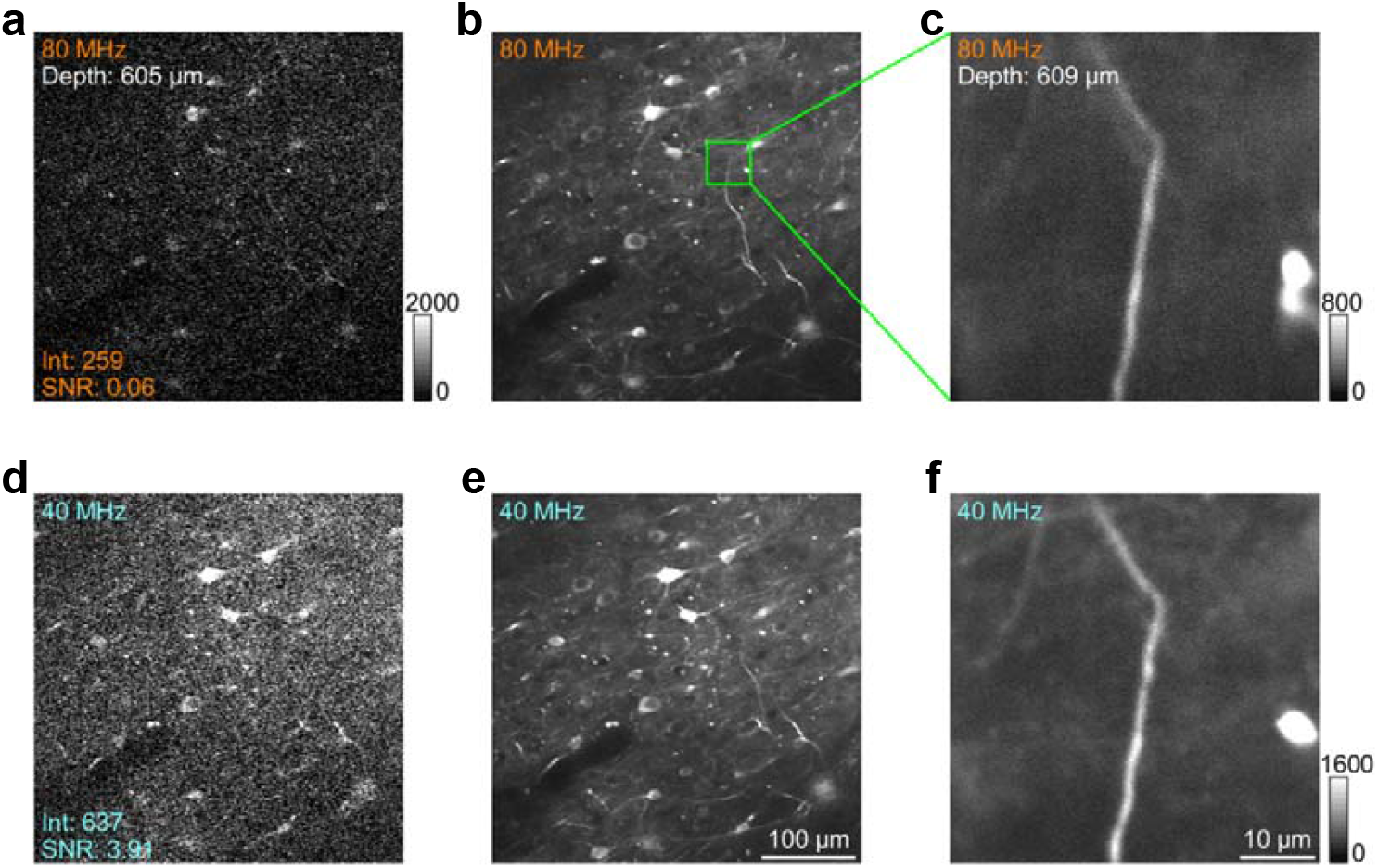
Superior deep-layer two-photon imaging of macaque visual cortex with 40 MHz laser excitation. All images were acquired at a depth of 605 μm and an average excitation power of 70 mW. **a** Raw two-photon image of neurons using standard 80 MHz laser excitation. **b** Averaged image from the 80 MHz dataset. **c** Zoomed-in image of a dendritic segment under 80 MHz excitation, showing limited structural detail. **d-f** Corresponding images using the 40 MHz frequency-divided laser: (**d**) raw image, (**e**) averaged image, and (**f**) zoomed-in dendrite image, demonstrating pronounced improvements in signal intensity, SNR, and structural clarity.

### 40 MHz repetition rate divider enhances spatial resolution in deep tissue imaging

At a constant average power, reducing the laser repetition rate from 80 to 40 MHz doubles the energy per pulse. Due to the nonlinear nature of two-photon absorption, this increased peak power is expected to not only boost fluorescence but also sharpen the effective point spread function (PSF), thereby improving spatial resolution. Our observation of a dramatic SNR increase in deep tissue strongly suggested this resolution enhancement. To quantify it, we measured the full width at half maximum (FWHM) of thin dendritic branches. Laterally, the 40-MHz configuration yielded significantly sharper images, with an average FWHM of 1.27 ± 0.152 μm compared to 1.35 ± 0.175 μm for the 80-MHz laser (p < 0.01, t-test; **Fig. 5a-d**). The improvement in axial resolution was even more striking: analysis of Z-series scans of dendrites revealed an axial FWHM of 5.2 μm for the 40-MHz mode, a marked improvement over the 9.3 μm measured for the 80-MHz mode (p < 0.001, t-test; **Fig. 5f, g**). These results confirm that the higher peak power at 40 MHz effectively sharpens the PSF, directly translating to a superior ability to resolve fine subcellular structures within deep scattering brain tissue.

**Fig. 5.**
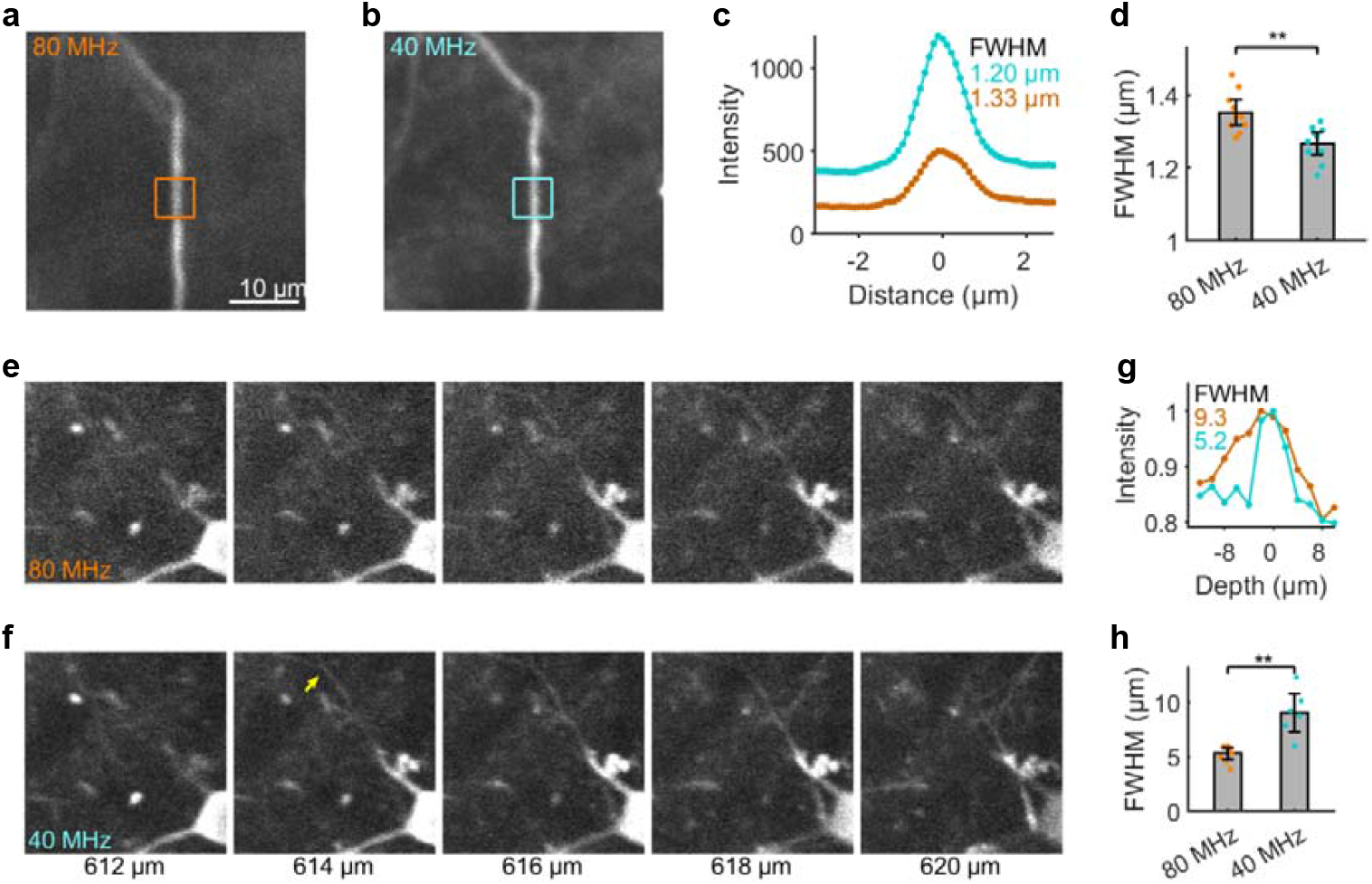
Lateral and axial resolution in deep-layer two-photon imaging using 80 MHz and 40 MHz femtosecond lasers. **a, b** Rotated dendritic images from **Fig. 4c** (80 MHz) and **4f** (40 MHz) for lateral resolution analysis; orange and light-blue squares denote regions of interest (ROIs). **c** X-axis intensity profiles of the ROIs (80 MHz: orange curve; 40 MHz: light-blue curve). Under 16 × zoom, 1 μm corresponds to ∼10 pixels. **d, e** Dendritic images from 5 consecutive z-planes (depth: 612– 620 μm); upper row: 80 MHz laser, lower row: 40 MHz laser. **f** Yellow arrow indicates the ROI for axial resolution measurement. **g** Axial intensity profiles (80 MHz: orange curve; 40 MHz: light-blue curve), normalized to the maximum intensity for each condition.

### Long-term stability and absence of detectable phototoxicity

A critical concern when increasing pulse peak power is the potential for phototoxicity. However, throughout our long-term imaging experiments, the fluorescence intensity of imaged neurons remained stable, with no evidence of photobleaching or tissue damage (Supplementary Fig. 6). This confirms that the pulse energies employed remain within a safe operational range for chronic cortical imaging. Furthermore, the frequency divider has been in continuous operation in our laboratory for over two years, demonstrating its robustness and long-term stability for demanding in vivo experiments.

## Discussion

In this study, we have developed and validated a straightforward and highly effective device for halving the repetition frequency of a standard femtosecond laser from 80 MHz to 40 MHz. By leveraging a Pockels cell in a resonant configuration, our system significantly enhances two-photon imaging performance, as evidenced by a two-fold increase in fluorescence intensity and more than a 3-dB improvement in SNR under identical average power conditions. These findings, demonstrated with both the widely used green calcium indicator GCaMP^6^ and the red fluorescent protein mScarlet3-H^17^, underscore the broad utility of this approach for diverse biological imaging applications.

The fundamental principle underlying this enhancement is the redistribution of laser energy. By doubling the time interval between pulses, the energy per pulse is doubled. Theoretically, this should result in a four-fold increase per pulse in two-photon excitation rate, thus around two-fold increase in two-photon imaging intensity because of half repetition rate of laser pulses. The robust signal enhancement observed is of significant practical value, especially for deep-tissue imaging or high-speed voltage imaging, which always suffer from low excitation efficiency and signal-to-noise ratios (SNR)^18-22^.

A critical consideration when increasing pulse peak power is the potential for accelerated photobleaching and increased phototoxicity^10,11,23^. While average power is linked to bulk sample heating^12^, high peak powers can induce higher-order nonlinear photodamage, leading to cellular dysfunction or death^11^. Our long-term stability experiments, which showed stable fluorescence over extended recording periods, suggest that at the power levels tested, our 40-MHz system does not induce acute phototoxicity and operates within a safe regime for longitudinal studies. Indeed, the enhanced efficiency allows for achieving a desired SNR at a lower average power, which can, in turn, reduce the overall photon dose and thermal load on the tissue, thereby minimizing phototoxicity and preserving sample health over time. This is a crucial advantage for chronic in vivo imaging, such as tracking neural plasticity or tumor progression over days or weeks^16^.

Compared to existing solutions, our frequency divider offers compelling advantages in terms of simplicity, cost, and ease of integration. Commercial pulse pickers and regenerative amplifiers are powerful but often complex and expensive, creating a high barrier to entry^13^. Our device, constructed from readily available components and custom electronics, can be implemented in any standard two-photon setup with minimal modification and expenditure. This accessibility empowers individual labs to upgrade their existing systems, making high-performance imaging more widely available. The resonant operation of the Pockels cell ensures high modulation efficiency and stability, contributing to the system’s robust performance.

In conclusion, we have presented a practical and powerful solution to a common limitation in two-photon microscopy. By efficiently converting an 80-MHz laser to 40 MHz, our frequency divider substantially improves imaging signal and quality without the complexity and cost of conventional methods. This work provides a valuable tool for the biological imaging community, enabling researchers to see deeper, faster, and longer into the complex world of living systems.

## Methods

### 40 MHz resonant frequency divider construction and operation

The frequency divider was engineered to halve the repetition rate of a standard 80-MHz femtosecond laser. For femtosecond lasers without direct 80 MHz signal output, a synchronizing 80 MHz signal could be obtained from the laser pulse via a commercial laser pulse detector (Thorlabs APD430A/M) or a simple photodiode circuity with a S13773 detector (**Supplementary Fig. 1**). A small laser energy was split onto the photodiodes using a 5% reflecting mirror or a simple 0.17 coverslip. This synchronizing signal was fed into an HMC432 frequency divider chip (**Supplementary Fig. 1)**, which generated a 40 MHz signal. This signal was then amplified to approximately 8 W using a 40 MHz, 25 W RF power amplifier powered by an adjustable DC supply (3–24 V). To drive the Pockels cell (Qioptiq LM 0202 P or Conoptics 350-80-02), the amplified RF signal was impedance-matched to a custom resonant LC circuit via a hand-wound 1:10 impedance transformer (**Supplementary Fig. 2, 3)**. The inductor (L1) was empirically optimized for each Pockels cell model: an 8-turn coil (8 mm outer diameter, 1 mm enameled copper wire) for the Qioptiq cell and a 12-turn coil for the Conoptics cell (**Supplementary Fig. 3)**. The resonant frequency was fine-tuned by adjusting coil tightness. The phase between the RF drive voltage and the arriving laser pulses was precisely matched by adjusting the length of the coaxial cable connecting the HMC432 output to the RF amplifier. In the correctly phased resonant state (**Supplementary Fig. 4)**, the Pockels cell functions as an electro-optic switch, routing every second pulse into a separate beam path and thus creating two independent 40 MHz pulse trains. The capacitance of the Pockels cell is temperature-dependent. To ensure both rapid start-up and stable operation, a constant temperature heating module maintains the cell at 31-32°C. We used a commercial temperature controller to drive a 10 W, 40°C PTC (Positive Temperature Coefficient) heater, which provides inherent safety against overheating (**Supplementary Fig. 3b)**. Additionally, because the L1 inductor carries a large resonant current when used with a Conoptics Pockels cell, a small cooling fin is required for thermal management.

### Animal subjects and surgical procedures

All experimental procedures involving two adult male rhesus macaques (Macaca mulatta, 5-7 years old, 5-8 kg) were performed in accordance with the guidelines for the Care and Use of Laboratory Animals of the National Institutes of Health and were approved by the Animal Care and Use Committee of Peking University.

To express fluorescent indicators, we performed viral injections into the primary visual cortex (V1). AAV9.hSyn.GCaMP5G^6^ or AAV9.hSyn.mScarlet3-H^17^ was pressure-injected (200 nL per site at 100 nL/min) at a depth of 300-500 µm below the dural surface. Following a 4-6 week incubation period, a chronic cranial window was implanted over V1. A custom-machined titanium head-post was affixed to the skull with dental acrylic to allow for head fixation during imaging sessions^16^.

### In vivo two-photon imaging

Imaging was performed on an Ultima IV (In Vivo) 2PM (Bruker Nano, FMBU, formerly Prairie Technologies). The excitation source was a Ti:Sapphire laser (Spectra-Physics, Mai Tai DeepSee) operating at a native repetition rate of 80 MHz, tuned to 1000 nm for GCaMP5G and 1040 nm for mScarlet3-H. The laser passed through the frequency divider before being directed to a scanning system composed of an 8 kHz resonant scanner and a galvanometer scanner. The beam was focused onto the sample through a water-immersion objective objective (16X, 0.8 N.A., Nikon). Emitted fluorescence was collected by the same objective, separated by a dichroic mirror, passed through appropriate emission filters (525/50 nm for GCaMP, 607/70 nm for mScarlet), and detected by GaAsP photomultiplier tubes (Hamamatsu, H7422-40). For all comparative experiments, the laser was switched between its native 80-MHz repetition rate (half power through a 50% transparent mirror) and a 40-MHz rate (using a frequency divider). The average laser power at the sample was kept constant for both conditions by adjusting a Pockels cell-based power controller (Conoptics) located behind the frequency divider. Images (512 × 512 pixels at zoom 2X) were acquired under resonant scanning mode at a frame rate of 30 Hz.

### Image analysis and quantification

All data were analyzed using custom scripts written in MATLAB.

### Image pre-processing

The images from each session were first realigned to a template image (an average image of 1000 frames) using a normalized cross-correlation-based translation algorithm, to correct the X-Y offset of images caused by the motion between the objective and the cortex^16^.

### Image quality

The Matlab function psnr was used to calculate the SNR of two-photon images (**Fig. 2** and **3**). An average template image (5000 frames for superficial layer imaging in Fig. 2 and 3, and 3,000 frames for deep layer imaging in **Fig. 4**) was used as reference images for psnr calculation.

### Spatial Resolution Estimation

To estimate effective spatial resolution, we analyzed high-magnification images of fine dendritic processes. The lateral resolution was determined by fitting a Gaussian function to the intensity profile of a cross-section perpendicular to a thin dendrite. The FWHM of the fitted Gaussian was taken as the lateral resolution. The axial resolution was estimated from Z-series stacks acquired with a 2-μm step size. The intensity profile of a horizontally oriented dendrite along the Z-axis was fitted with a Gaussian, and its FWHM was taken as the axial resolution.

## Statistical Analysis

Comparisons between 80-MHz and 40-MHz conditions were performed using a two-sided paired t-test. Differences were considered statistically significant at p < 0.01. Data are presented as mean ± s.d. unless otherwise noted.

## Data availability

The data that support the plots within this paper and other findings of this study are available from the corresponding authors upon reasonable request.

## Code availability

The code that support the plots within this paper and other findings of this study are available from the corresponding authors upon reasonable request.

## ACKNOWLEDGMENTS

We thank Jianwei Zong and Ian Andolina for help in manuscript revision, Yang Li and Hongfei Jiang for assistance in animal preparation. This work was supported by National Natural Science Foundation of China (grant no. 31730109 and U1909205) and funds from the Peking-Tsinghua Center for Life Sciences.

---

**Supplementary Fig. 1.**
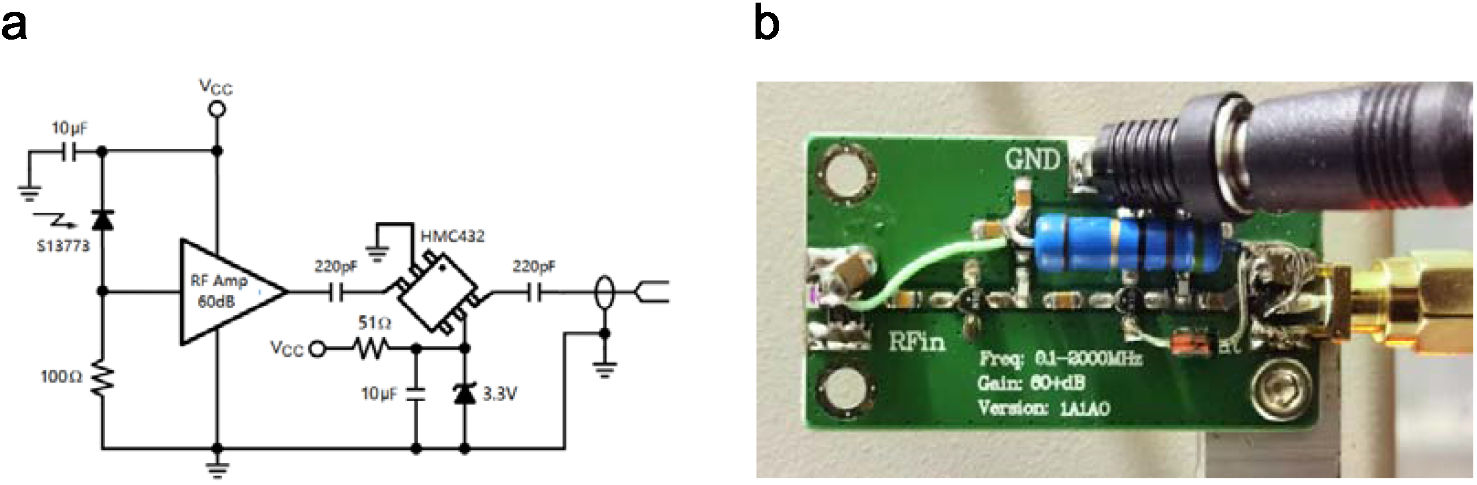
Integrated Laser Pulse Detection and Frequency Division Module. **a** Schematic of the electronic circuitry for an integrated module designed to detect 80 MHz laser pulses and perform frequency division. **b** A demonstration of an integrated RF amplifier board with photodiodes and frequency divider.

**Supplementary Fig. 2.**
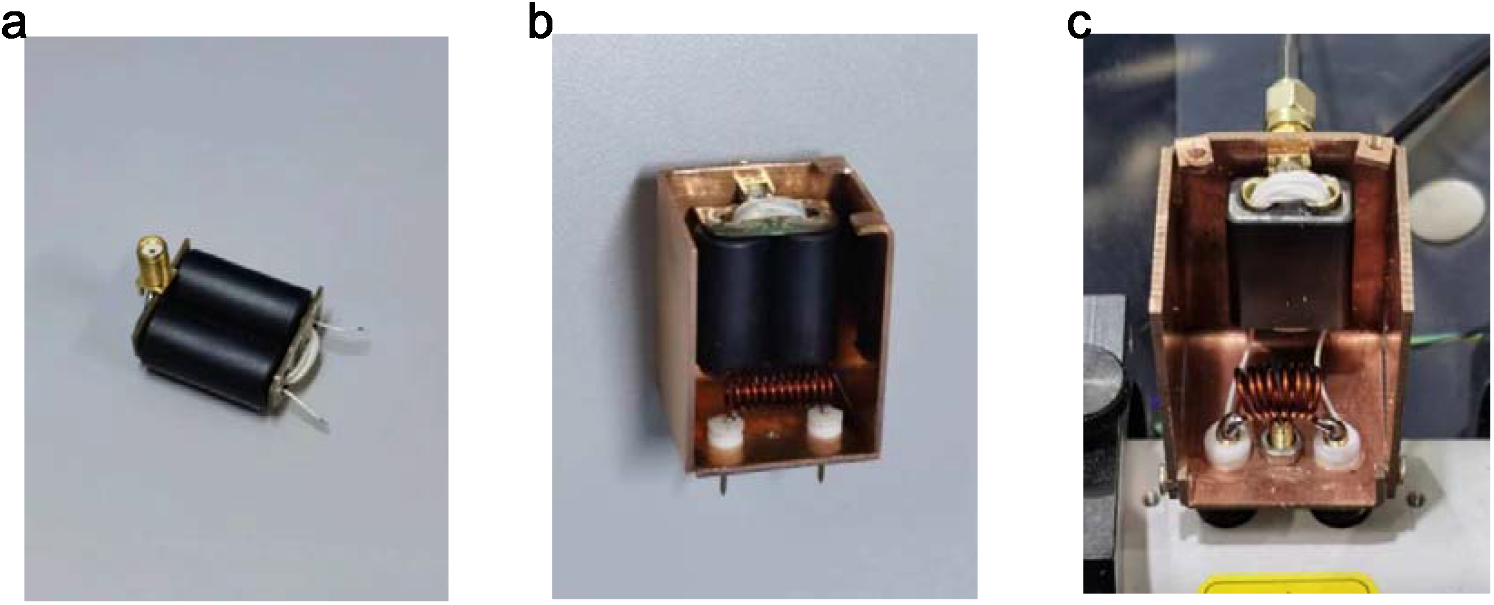
Photographs of Impedance Transformer B1 and Resonant Inductor L1. **a**-**c** Assembly of the impedance transformer (B1) and resonant inductor (L1) within a copper shielding box.

**Supplementary Fig. 3.**
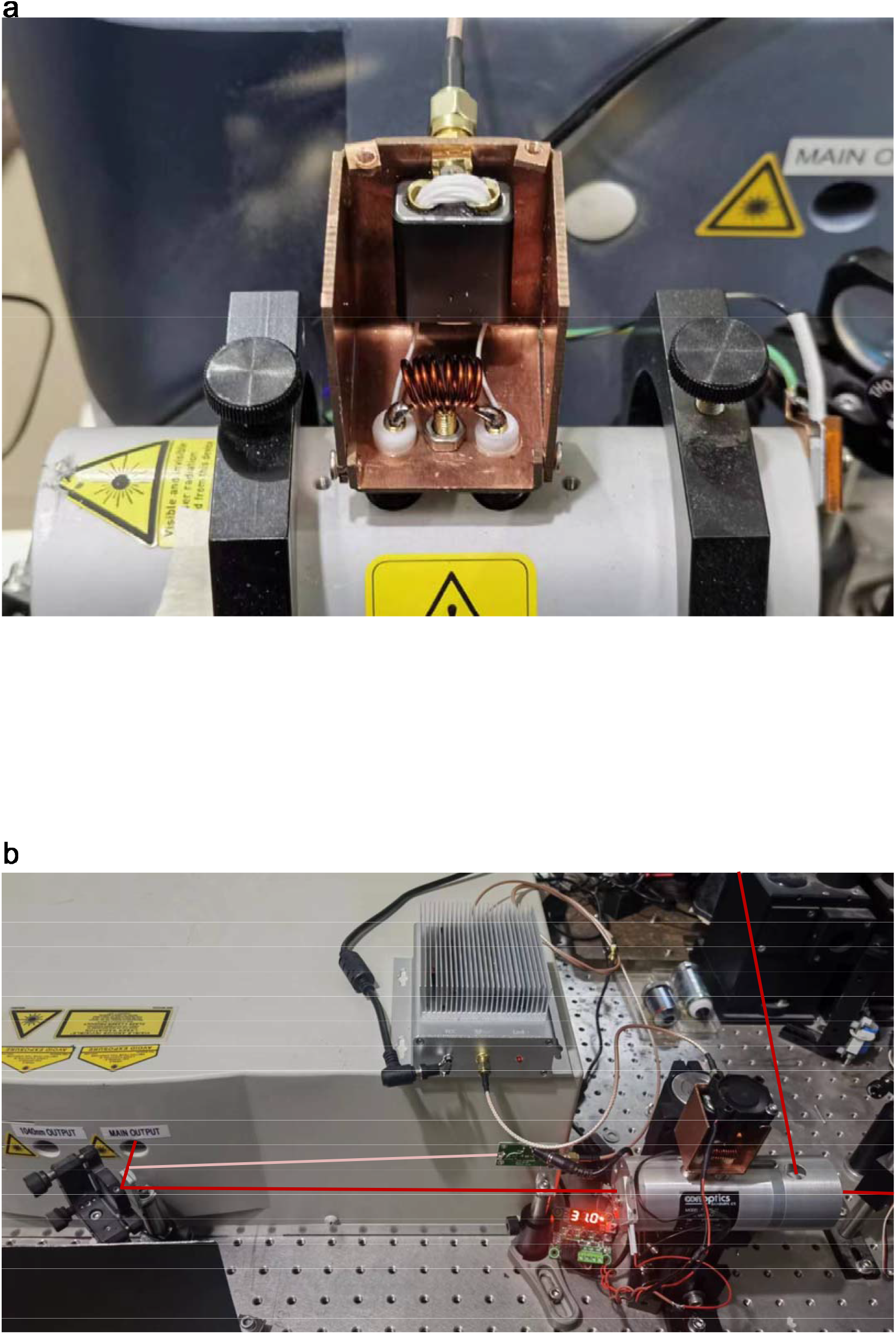
Demonstration of a 40 MHz Femtosecond Laser Repetition Frequency Divider. **a** RF transformer unit connected to a Qioptiq LM 0202 P Pockels cell. **b** Photograph of a laser repetition frequency divider using a Conoptics Model 350-80-02 Pockels cell. A 0.17 mm cover glass was employed to reflect ∼5% of the power from an InSight DeepSee femtosecond laser onto a photodiode, generating an 80 MHz synchronization signal. The pink line denotes the ∼5% laser power to the photodiode; the dark red lines indicate the main laser power paths.

**Supplementary Fig. 4.**
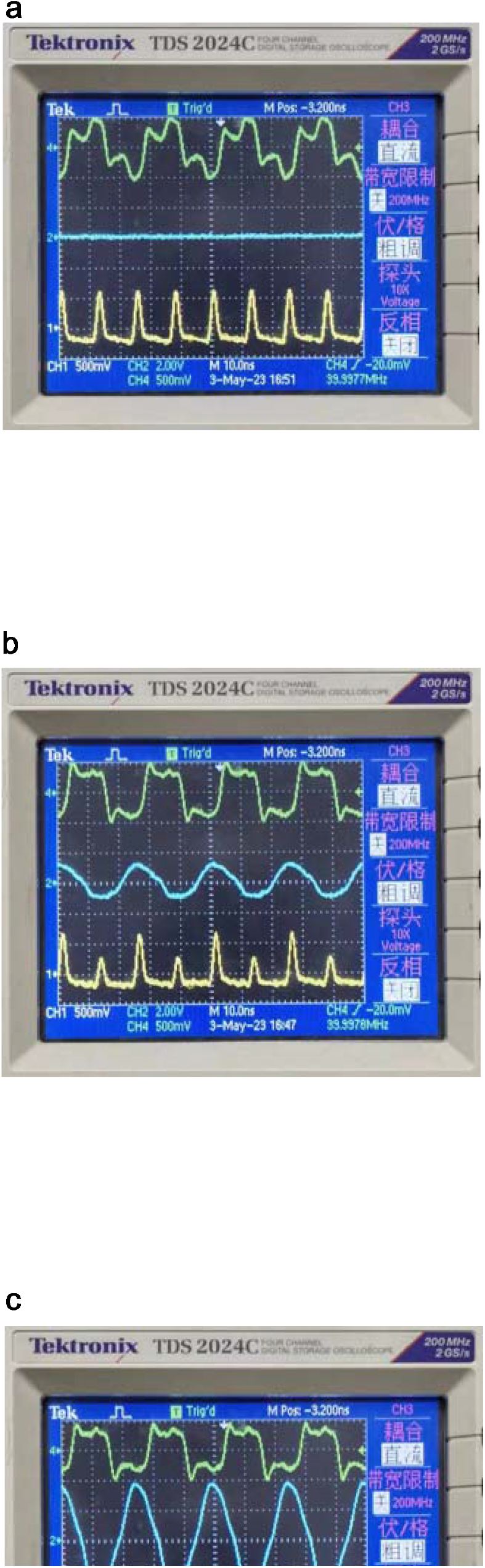
Electronic Signals of a Demonstration Femtosecond Repetition Frequency Divider under Varied RF Power Amplifier Supply Voltages. The green trace (CH4) represents the 40 MHz synchronization signal; the blue trace (CH2) represents the inductive signal from the resonant circuit; the yellow trace (CH1) represents the output laser pulse signal detected by an additional photodetector. **a** When the RF power amplifier supply voltage is set very low (0–5 V DC), no pulse modulation is observed. The output laser pulses maintain an 80 MHz repetition rate because no 40 MHz resonant voltage is applied to the Pockels cell. **b** As the supply voltage is increased to 10 V, the Pockels cell partially modulated the laser pulses. **c** When the supply voltage is further increased to 18 V, the Pockels cell completely blocks the interval laser pulses (reflected to direction B, as shown in **Fig. 1a**), resulting in the output of a 40 MHz femtosecond laser pulse train.

**Supplementary Fig. 5.**
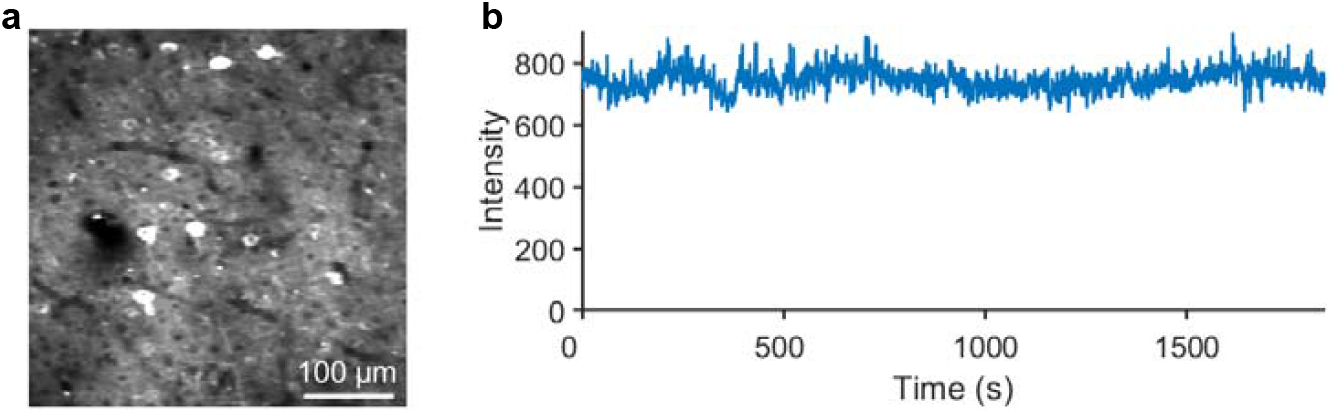
Long-term stability of two-photon imaging using a 40-MHz femtosecond laser. **a** Two-photon image of GCaMP5G-expressing neurons in the macaque visual cortex, acquired using a 40-MHz femtosecond laser at an average power of 40 mW. **b** Plot of the average fluorescence intensity from the field of view over a continuous 30-minute recording period, demonstrating stable signal levels.

